# *Live-action* active methods for biochemistry and pharmacology learning

**DOI:** 10.1101/2025.06.10.658848

**Authors:** Marco Augusto Sperandeo Liguori, André Ferreira Aparício, Thomaz A. A. Rocha e Silva

**Affiliations:** Academic, Faculdade Israelita de Ciências da Saúde Albert Einstein, Hospital Israelita Albert Einstein, São Paulo, Brazil; Faculdade Israelita de Ciências da Saúde Albert Einstein, Hospital Israelita Albert Einstein, São Paulo, Brazil

**Keywords:** Live-action active method, pharmacology teaching, biochemistry teaching

## Abstract

The integration of active learning strategies in health education is essential for improving student engagement and understanding, particularly in basic sciences like pharmacology and biochemistry. However, these disciplines are often perceived as abstract and disconnected from clinical practice. The aim was to develop and evaluate two innovative, live-action simulations for teaching fundamental concepts in pharmacokinetics and enzyme kinetics, using accessible materials and student-centered design. The study involved 305 students from health-related undergraduate and graduate programs. Two simulations were designed: a pharmacokinetics (PK) simulation representing systemic circulation and drug metabolism using plastic blocks and classroom stations; an enzyme kinetics (EK) simulation using both digital slides and physical block sets to model substrate-product reactions and construct Michaelis-Menten curves. Student perceptions were measured using a Likert-scale instrument adapted from validated educational models. Quantitative data (pharmacokinetic and enzymatic curves) were also generated from simulation outcomes. Students successfully built representative pharmacokinetic and enzymatic activity curves, allowing exploration of key concepts such as drug absorption, metabolism, Cmax, Tmax, half-life, and enzyme-substrate reaction rates. Perception surveys revealed high approval levels, particularly regarding the effectiveness, engagement, and collaborative aspects of the simulations. Over 85% of participants preferred the simulation-based approach over traditional lectures. These live-action activities represent a novel, effective, and low-cost strategy for enhancing the teaching of basic sciences in health education. The methods promote experiential learning, integrate cognitive, affective, and psychomotor domains, and foster student autonomy and motivation. These findings support the broader application of active methodologies in foundational science curricula.

## INTRODUCTION

Basic sciences are the foundations of any professional educational system and generally offered as difficult disciplines in the beginning of graduation programs. In health professions, biochemistry is usually one of the first basic sciences disciplines to attend besides anatomy and cell biology. Otherwise, pharmacology is thoughtfully positioned as one of the last disciplines of basic cycle, due its prerequisites of other basic disciplines.

The effectiveness of various pedagogical approaches in biochemical and pharmacological education has been the focus of recent research, highlighting the need for innovative strategies to enhance student engagement and comprehension. Active learning techniques, such as problem-based learning, interactive lectures, and team teaching, have demonstrated significant improvements in student retention and conceptual understanding (Govindarajan et al., 2021)

The integration of interdisciplinary approaches, particularly in pharmacology education, is essential to bridge the gap between theoretical knowledge and clinical application. Studies conducted in Brazilian medical schools have indicated that while traditional expository lectures remain the predominant teaching method, there is an increasing recognition of the need for more dynamic and student-centered strategies to enhance motivation and performance (Fidalgo-Neto et al., 2023)

Recent advances in pharmacology and biochemistry education emphasize the importance of active learning strategies and pedagogical content knowledge to enhance student engagement and comprehension. Engels (2018) discusses the necessity of rethinking pharmacology curricula to integrate competency-based learning approaches that align foundational pharmacological principles with clinical application. Despite the widespread adoption of integrated curricula, concerns remain regarding the adequacy of pharmacological knowledge among graduates, as insufficient training has been linked to prescribing errors and medication-related adverse events. To address these gaps, educators are exploring active learning techniques such as case-based learning, problem-solving exercises, and interactive pedagogical models. Similarly, Kopecki-Fjetland and Steffenson (2021) advocate for an iterative approach to active learning in biochemistry education, highlighting the effectiveness of diverse methodologies, including problem-based worksheets, tactile learning activities, and learning cycles, in reinforcing core biochemical concepts. Both studies underscore the critical role of student-centered, evidence-based teaching strategies in fostering deeper learning and long-term retention, suggesting that the incorporation of active learning frameworks could significantly improve pharmacological and biochemical education outcomes.

To address these challenges, various simulation models have been developed to support clinical and bedside learning, while basic sciences are often relegated to digital models as a comprehensive platform for understanding molecular and cellular processes. In this study, we designed two live-action simulations for biochemistry and pharmacology education, translating molecular-level phenomena into hands-on, manipulable activities, thereby giving practical meaning to the *live-action active methods*.

## MATERIAL AND METHODS

### 1. Local of research

Research was conducted in the Center of Education and Research of the Albert Einstein School of Health Sciences, São Paulo, Brazil. This Faculty is linked to Albert Einstein Israelite Hospital, São Paulo, Brazil. The two sections of the project (as described as follows) were separately registered under institutional numbers 4001-19 (pharmacokinetic) and 4679-21 (enzymatic kinetic).

### 2. Sections of the project

The project was divided in two sections for bureaucratic and regulatory purposes. The first section was the pharmacokinetic and the second was the enzymatic kinetic section. Although conducted separately, they shared the main core of transposing molecular phenomena to a metric-based simulation, including resources, challenges, limitations and outcomes. Therefore, those sections will be referred as PK and EK for pharmacokinetic and enzyme kinetics respectively along the manuscript.

### 3. Research population

A total of 305 students were enrolled in research, with 256 in PK and 49 in the EK. In the PK section, students from Dentistry, Medical and Nursing school and Neuroscience Post graduation participated, with 94 answering the perception test. In the EK section, all students performed the simulation and answered the perception scale.

#### 3.1. Inclusion criteria

Undergraduate and post-graduation health students performing disciplines correspondent to project sections, i.e., biochemistry for EK and pharmacology for PK. Students of Medicine, Nurse, Physiotherapy and Dentistry graduation courses were included, also Neurosciences Post-graduation.

#### 3.2. Exclusion criteria

Students who did not consent to volunteer and classes that did not follow the instructions of the simulation.

### 4. Ethics

Every volunteer was submitted to the Free Informed Consent process and signed the Consent Term online (PK) or physically (EK). Protocols were submitted and approved by Brazilian Research Ethics System, as stated in decisions 6.108.603 (PK) and 5.156.641 (EK), available in https://plataformabrasil.saude.gov.br/login.jsf

### 5. Methods

#### 5.1. Pharmacokinetic simulation

The objectives of this simulation were to build two pharmacokinetic curves, compare the curves of two different ways of administration and determine pharmacokinetic parameters in the curves. It is mandatory that students had read a previous synthetic material on molecular processes of pharmacokinetic: administration, absorption, distribution, metabolism and excretion. The simulation is conducted by the teacher, organizing students’ dynamics.

It consists of setting the classroom into a human body circulation model (appendix A) with defined stations in tables and students performing the blood circulation. Stations are brain, right heart, lungs, left heart, intestine, liver, kidney, peripheral organs and vein access. It is important that stations disposition and circulation circuit reproduce basic human anatomy, such as an exclusive circulation to the brain, a direct circulation from intestine towards liver, and independent circulation to kidney and peripheral organs which converge to a common access to right heart, sided by the vein access (appendix A). Also, five students are required to specific tasks: two in liver, one in intestine, one in kidney and one in time and data control.

The drug is represented by a two pieces plastic block, from commercial brand. Once the blocks are inserted in simulation (administered), students will carry them in circulation circuit, leaving OR catching in each station, one block per hand. Students in the liver station has to modify the two-block to a three-block drug, inserting a small block into structure, simulating Phase 1 metabolism. Also, they must transform the three-block to a four-block “drug”, connecting a larger block to simulate Phase 2 conjugation. The student in kidney removes only the four-block drug left in the station.

Two rounds were performed: intravenous and oral administration. For intravenous, the drugs are left in vein station and passing students collect it to transport. For oral administration, a pot with drug is left with the student in intestine who constantly puts the blocks one-by-one in the station.

It is mandatory that students perform aleatory circulation and blocks managing, i.e., they must not choose where to go, walking always and only to empty spaces, and do not choose which blocks to catch, acting blind when picking a piece.

Finally, the controller student pauses the simulation in predetermined intervals with a STOP oral command and counts the active (two-block) drugs in students hands. The result of each interval is used to build the pharmacokinetic curve.

#### 5.2. Enzyme kinetic simulation

The objectives of this simulation were to build the enzymatic activity curve, note the reaction equilibrium moment and extrapolate data to Michaelis Menten curve construction, while students play the enzyme role. Two methods were applied aiming the optimization of the simulation in the classroom context. The first was an online-based simulation in which a cloud presentation archive is shared with the students (appendix B). Each archive has six slides crowded with small black spheres, but each slide has also pairs of semitransparent yellow and blue spheres, in the role of the substrates. The first slide has two pairs, the second has four, and eight, 16, 32 and 64 sequentially towards the sixth slide. The goal is to manually merge blue and yellow spheres to a green one, then called product. Each group of seven students has one archive and each student manages only one slide of it. The teacher starts chronometer saying “GO!” and students starts merging the blue and yellow spheres. At predetermined intervals, the dynamic is paused and the cumulative number of green spheres is registered. The simulation may last until the curves start flattening.

The second method used plastic blocks. Black and white blocks filled a pot and pairs of blue and grey blocks were added to five pots (appendix B). The first had five pairs, the second 10, and 15, 20 and 25 sequentially to the fifth pot (see Figure 3). Each group of students had the five pots and each student managed one pot, sided to a stopwatch controller student to each one. The simulation consists of finding and connecting blue and grey blocks, forming the product. The start is the stopwatch trigger and each product made is registered as a lap. At the end of substrate, the time is considered cumulative at each product formation.

#### 5.3. Students’ perception

The perception of the students about each simulation was measured using a scale adapted from the work of Eukel *et al.*, (2017) in previous works (Cunha *et al.*, 2023), with 10 statements subdivided into three dimensions: support for learning, learning activity, and collaborative learning. The instrument was adapted by the authors specifically for the present work, as follows.

S1 - The virtual escape room encouraged me to think about the material in a new way S2 - I would recommend this activity to other students

S3 - I learned from my colleagues during the virtual room

S4 - The virtual room is an effective way to review the topic of AMI, Nursing care and drugs used to treat patients with this condition

S5 - The virtual escape room is an effective way to learn information related to the clinical conditions of AMI patients

S6 - I learn better with the game format than the lecture

S7 - I feel that I was able to engage with my teammates to learn the material

S8 - It was difficult for me to focus on learning because I was feeling stressed or overwhelmed

S9 - Non-educational parts (e.g. crossword puzzles) distracted me from learning about clinical conditions of AMI patients

S10 - I prefer to gather information from multiple sources for my learning

For each statement an answer scale was defined as: I strongly disagree with the statement; I disagree with the statement; I neither agree or disagree with the statement; I agree with the statement; I strongly agree with the statement.

## RESULTS

### 1. Pharmacokinetic simulation

The construction of a didactic pharmacokinetic graphic was achieved in different classroom configurations, considering number of participants and proportion of “drugs administered”. Classes with 28, 43 and 62 students created similar graphs which represented intravenous and oral administrations of drugs (Figure 1 A, B and C). More, the proportion of one “drug” for each student in “intravenous” administration, and the double in “oral”, were suitable to build representative curves which allow exploring concepts such as absorption rate and first pass metabolism. The concepts of Cmax, Tmax, half-life and Area Under Curve were clearly explorable.

**Figure 1.**
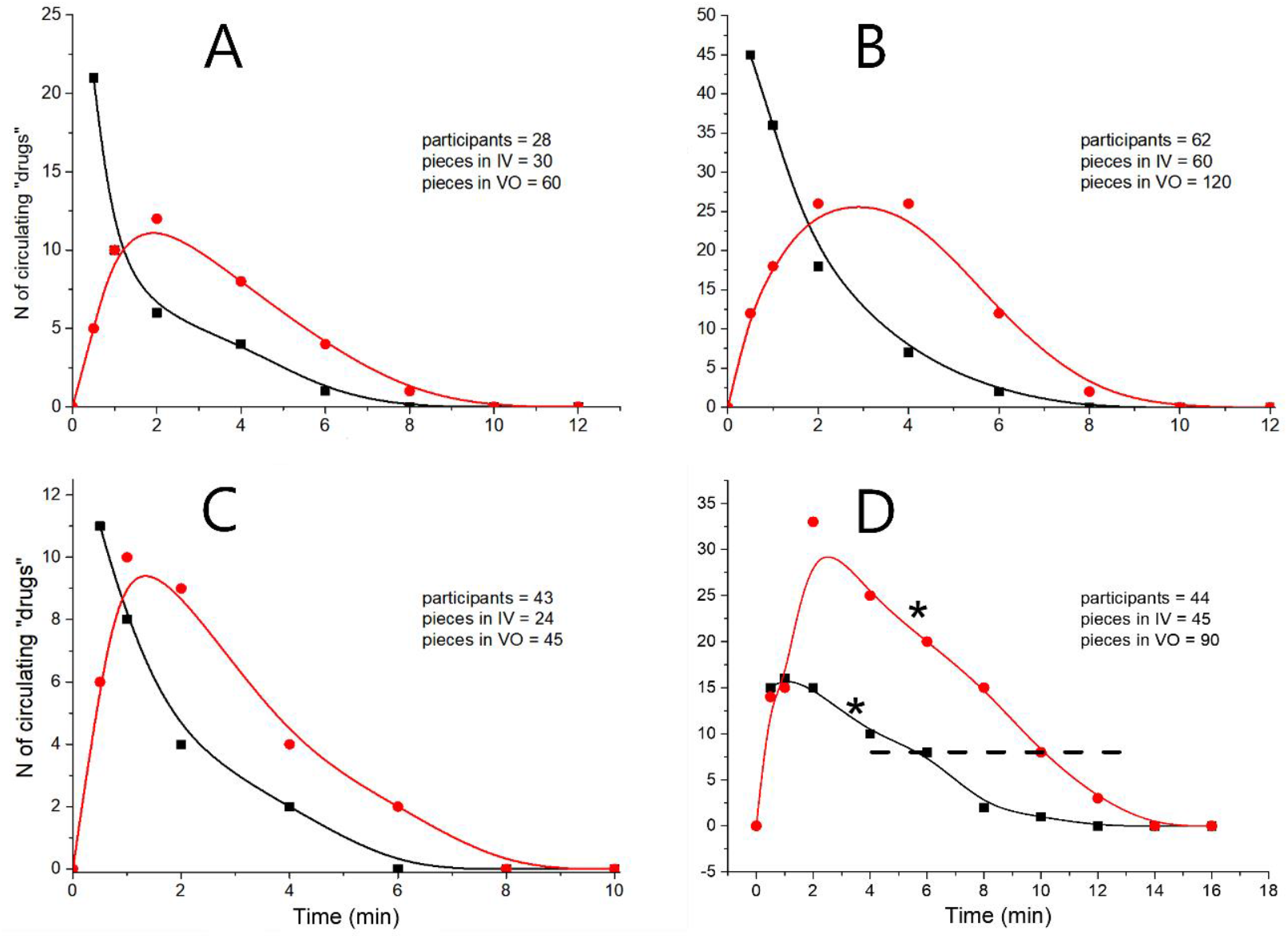
Pharmacokinetic curves obtained with different number of participants and proportions of “drugs” (A, B and C). (D) Pharmacokinetic curve obtained in a session which students in liver station performed a slow metabolism. Note the zero order phase of the curve (*) and regular first order decay after critical concentration lowering (below dashed line). Black curves: IV – represents the method that simulates intravenous administration; red curves: VO – represents the method that simulates oral administration.

Four representative curves obtained are presented in Figure 1. Accidentally, one class had students in liver station who did not performed “drug transformation” in a maximum speed, and a zero order metabolism pharmacokinetic curve was obtained (Figure 1D).

The students’ perceptions of pharmacokinetic simulation were obtained from 94 participants described in table 1. It was positive in all aspects, with statements 1 (The simulation encouraged me to think about material in a different way) and 4 (The simulation is an effective way to review the topic of pharmacokinetics) with highest agreement levels. The negative (S8 and S9) and neutral (S10) statements had the most diverse scores, but still with over 75 and 65% of approval, respectively (Figure 2).

**Table 1.** Number of participants of the pharmacokinetic simulation and respondents of the respective perception scale.

| Course | Simulation | Perception scale |
| --- | --- | --- |
| Dentistry | 43 | 0 |
| Medicine | 101 | 32 |
| Nursing | 68 | 45 |
| Physiotherapy | 28 | 0 |
| Neuroscience (post-graduation) | 16 | 17 |
| Total | 256 | 94 |

**Figure 2.**
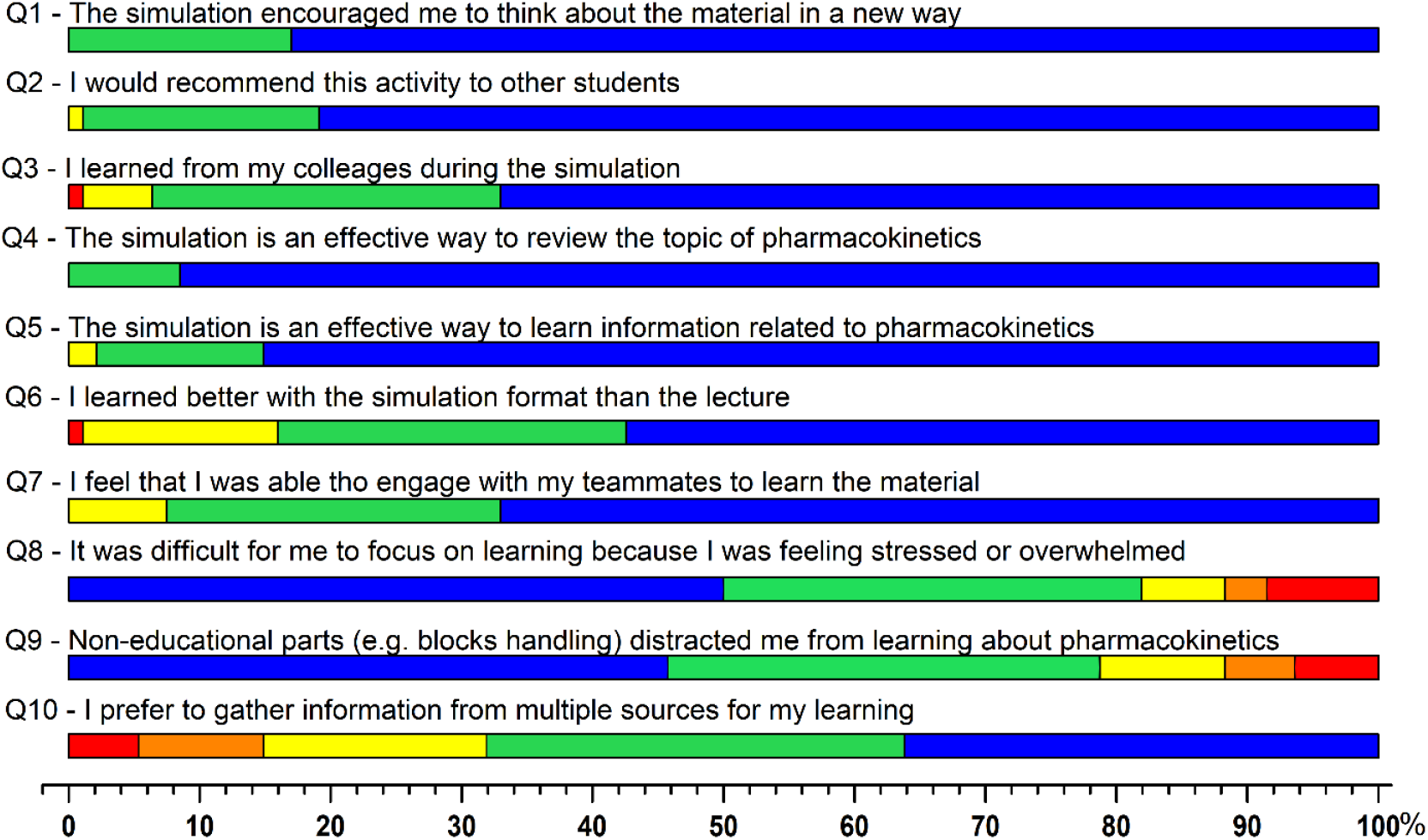
Perceptions of the students about pharmacokinetic simulation (N = 256). Colors were attributed to answers in a spectral distribution: I strongly disagree with the statement – red; I disagree with the statement – orange; I neither agree or disagree with the statement – yellow; I agree with the statement – green; I strongly agree with the statement – blue. Note that for negative statements (S8 and S9), colors spectra were inverted.

### 2. Enzymatic kinetic simulation

The simulations were performed in digital and physical methods as detailed in Methods section. The product formation graphics obtained in both methods are shown in figure 3.

**Figure 3.**
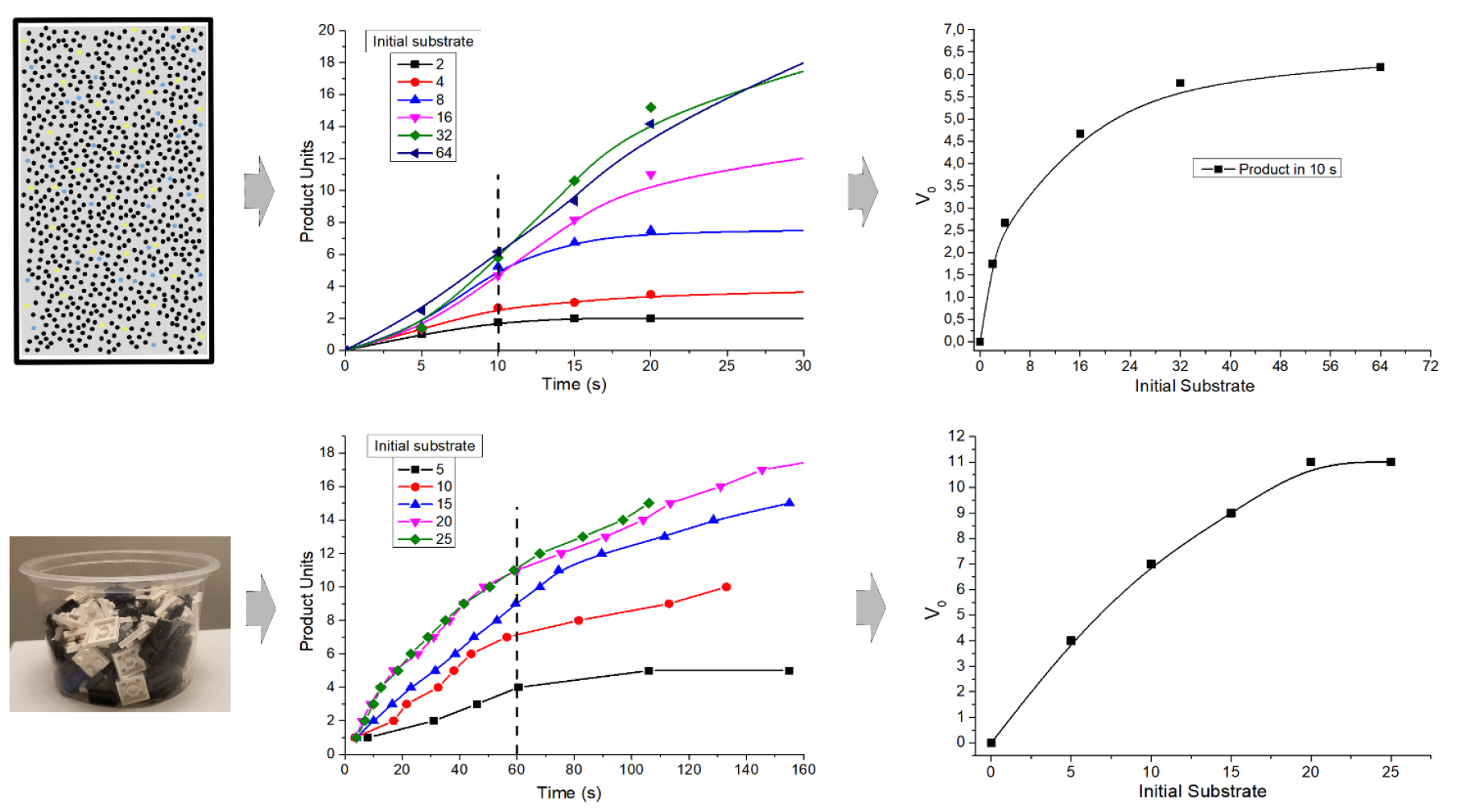
Enzymatic kinetics simulation using digital (up) and physical (down) methods. The figures in right illustrates the source of each method, the graphics in the middle are product formation curves and left are the Michaelis Menten graphs extrapolation of the curves in times indicated by dashed lines.

**Figure 4.**
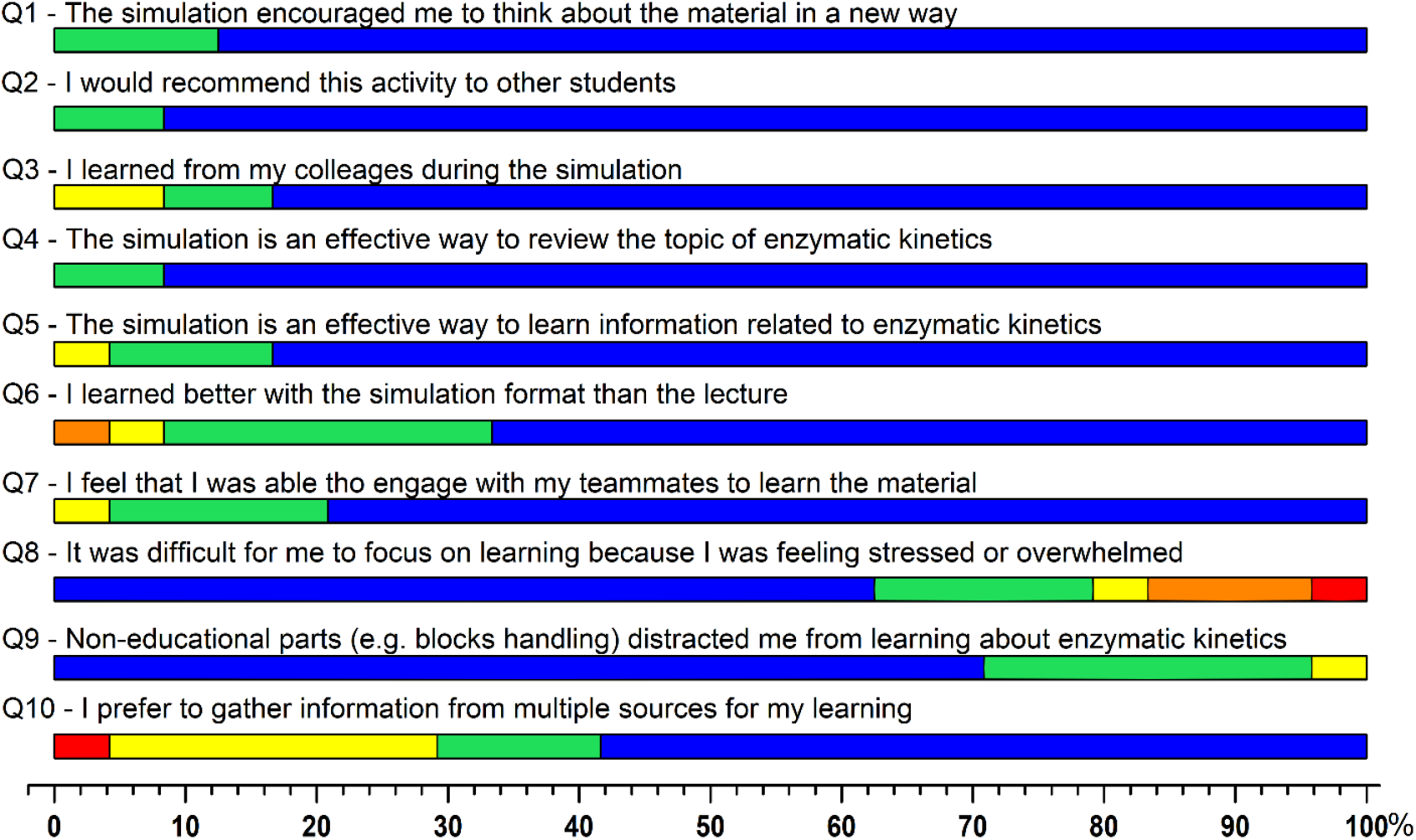
Perceptions of the students about pharmacokinetic simulation (N = 49). Colors were attributed to answers in a spectral distribution: I strongly disagree with the statement – red; I disagree with the statement – orange; I neither agree or disagree with the statement – yellow; I agree with the statement – green; I strongly agree with the statement – blue. Note that for negative statements (S8 and S9), colors spectra were inverted.

The digital method provided a more organized graph, once the time for product count was equal to every participant. On the other hand, the plastic blocks method showed a more realistic curve for product formation. In both cases the product curves were successfully build. The initial equilibrium was evident while students reached a constant rate of “product” synthesis, what was reinforced with the growing difficulty to perform the reaction with substrate rarefaction. The extrapolation for Michaelis Menten curve was possible and the concept of substrate dependent rate of reaction (V_0_) became clear.

The perception of the students was positive with three statements reaching 100% of agreement: The simulation encouraged me to think about material in a different way; I would recommend this activity to other students; The simulation is an effective way to review the topic of pharmacokinetics. Also, the negative statements had a disagreement over 75 and 95% (S8 and S9, respectively). Figure four shows the score of students’ perceptions for enzymatic kinetics.

## DISCUSSION

The term *live-action* was applied to this work aiming to define the transposing of a molecular phenomena from an angstrom and molar scale to the hand-made metric dimension. Despite the challenges of such adaptation, several digital models and platforms were successfully developed for molecular sciences teaching (Florjanczyk *et al.*, 2018; Mak *et al.*, 2024). In other hand, simulation is a potent tool often conducted in digital environment or actor-driven sessions for teaching applied and clinical heath sciences (Khan *et al.*, 2024).

This work brought to live-action two cornerstones of heath professional curricula, offering to students a dynamic interactive activity that allows the direct observation of the phenomena. The authors opted for applying a perception scale rather than adopting a test based experiment due to the bias in quasi-experimental or the privation of activity in control groups in experimental design. More, the live-action ought the students to bring previous knowledgement and apply it (cognitive domain) in a group coordinated and hermetic ruled dynamic (affective domain), while perform physical exercise and interactions (psychomotor domain). Therefore, the results shows that students actually liked the simulation for several reasons: new way of thinking stimulation, collaborative learning, information revision and learning, non-educational activities during learning. The limitations of each simulation are discussed as follows.

The pharmacokinetic simulation presented curves similar to those in clinical studies (Hernández-Jiménez *et al.*, 2024; Fiori *et al.*, 2025), representing a solid source for curve construction comprehension and parameters settings. During the simulation, the STOP command calls student’s attention to “drug” counting and then, the presentation of the final curve, allows participants to feel enrolled since it was built with the data they generated. Parameters as steady state concentration (Css or Cmax), maximum time (Tmax), area under curve (AUC) and half-life are fully understandable and comparable in two modes of “administration”. Also, half-life concept, which it is a property of the drug while in blood independent of the route of administration, was demonstrable too. More, during (or even after) the simulation the teacher can point to processes as: drug absorption in gut and first-pass metabolism influence in a doubled dose oral administration; biochemical function integration between liver and kidney; simultaneity of absorption, distribution, metabolism and excretion and aleatory circulation of active drug and its metabolites. Nevertheless, if the teacher keeps those topics and gives room to student’s considerations immediately after the simulation, they shall reach those concepts by themselves, only guided by the teacher’s tips and instigating questions.

The limitations of pharmacokinetics simulation are divided into mechanistic and logistic aspects. The mechanistic is inherent of the live-action feature, since the number of drug blocks is proportional of “blood transporters”. It does not allow to explore concepts like free and protein-conjugated drug, or the tendence to zero in final depuration – since our model indeed reach zero. The logistic issue requires a tight conduction of the teacher, giving clear and repeated instructions to participants, reinforcing mandatory behaviors such as not choosing where to go our drug blocks to catch. It is also difficult motivating the movement during this repeated physical activity in an each time more sedentary student population. Although, the simulation showed to be consistent and plausible to be explored, even with slight deviations that reflected in curve distortions. An example was the zero-order pharmacokinetic curve that was unpredictably obtained when students in the liver station were dispirited and did not perform the block changing at their maximum speed.

Other active methods published in pharmacology learning includes computer and case-based techniques, flipped classroom, modules and others, with a test score based experimental design (for review, see Jaju *et al.*, 2024). Although, Granat *et al.* (2024) explored gamification learning with pre and post-test in an experimental design, demonstrating the effectiveness of such approach. More, their work associated a “yes or no” answered survey about students’ impressions of the educational game with almost unanimous approval. This demonstrated the relation between students’ perception and test grade, explored ahead.

The enzyme simulation enabled students to grasp a fundamental concept of (bio)chemical reactions: the concentration-dependent nature of favorable molecular collisions. Since participants were required to pair a yellow sphere with a blue one to form a product, the difficulty of this task was inversely proportional to the concentration of available spheres. This experience led students to intuitively understand that enzyme activity is dependent on substrate concentration. By observing their peers, students could compare product formation rates and perceive the decreasing speed of substrate discovery and reaction over time. These dynamics were reflected in the product formation curves. Subsequently, by deriving the initial reaction rates and plotting them against substrate concentration, students were able to construct the Michaelis-Menten curve using their own data.

The main limitation of the live action enzyme simulation lies in the inability to translate molar particle quantities into real-life physical interactions. A key biochemical principle—namely, that reaction equilibrium occurs when substrate concentration vastly exceeds enzyme concentration—cannot be fully modeled in this format. Nonetheless, the simulations produced linear initial phases, aligning with theoretical expectations. Variations in student skill levels with digital tools or physical materials also influenced outcomes, but this could be minimized by having multiple students perform the same task and averaging the results.

Regarding the digital method, device type proved to be a limiting factor. Smartphones performed worst, followed by tablets. The most effective devices were laptops with mouse input, which facilitated better control and faster interaction.

As for the perception scale, students’ responses reaffirm the suitability of active methodologies in basic science education. For both simulations, statements related to engagement, topic review, and learning outcomes received overwhelmingly positive responses, while negative statements showed no more than 20% agreement or neutrality. Factors such as individual stress or distraction from non-educational elements are inherent challenges of active learning methods and are expected in any voluntary experimental group (Cunha et al., 2023). Furthermore, although there was strong agreement (at least 85%) with the statement “I learned better with the simulation method than with traditional lectures,” a notable portion of students still preferred conventional formats. This suggests that a balanced integration of active and traditional teaching approaches may represent the most effective strategy in health education.

In conclusion, this study introduced two innovative active learning strategies—based on live-action active methods—for teaching pharmacokinetics and enzyme kinetics. These simulations received strong approval from students and expanded the repertoire of accessible, interactive tools for teaching fundamental scientific concepts in health education, reinforcing core theoretical principles through practical, hands-on learning.

## AKNOWLEDGEMENTS

The original ideas of the present project were generated during the 2020 International Scholars Collaboration in Teaching and Learning program of the Case Western Reserve University, Cleveland, USA. The authors would like to thank the first class in Frontiers of Neuroscience Post-graduation Program for pilot study collaboration. The authors also thank all undergraduate students in Medicine, Nursing, Physiotherapy and Dentistry who volunteered to this research. This work was supported by Fundação de Amparo à Pesquisa do Estado de São Paulo – FAPESP process number 2021/04004-7.

## APPENDICES

### Appendix A

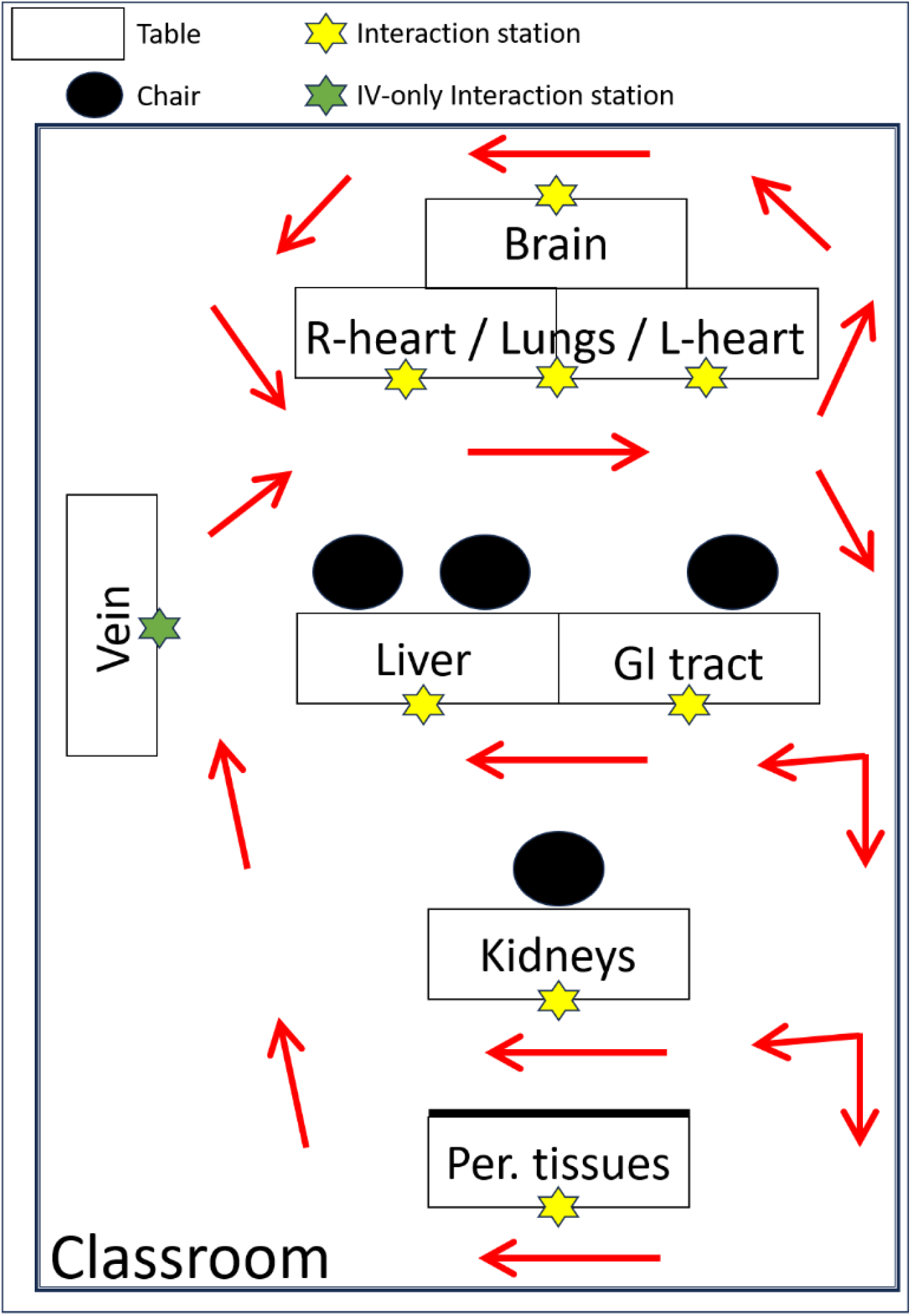

### Appendix B

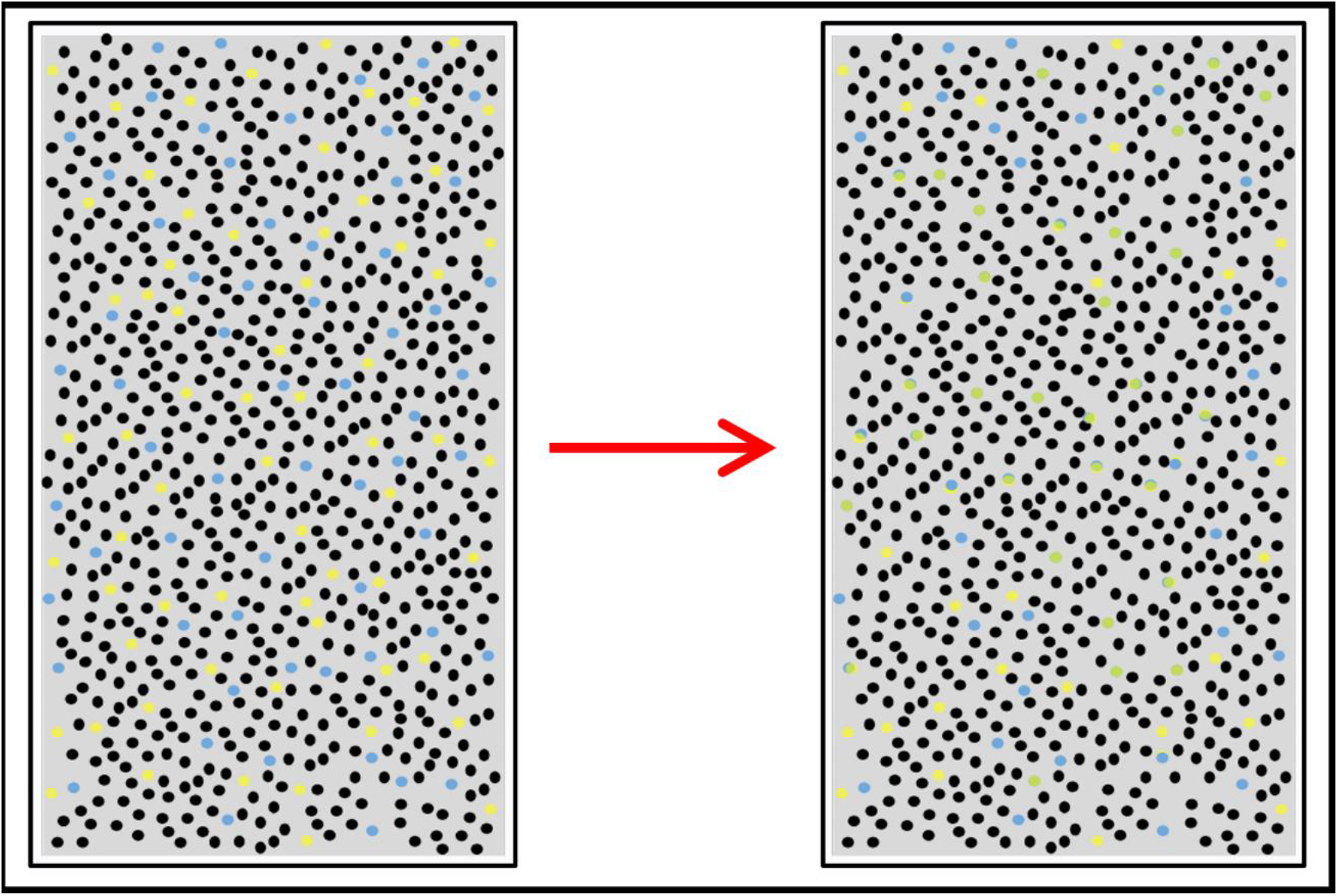

